# Exploring ligands that target von Willebrand factor selectively under oxidizing conditions through docking and molecular dynamics simulations

**DOI:** 10.1101/2024.03.22.586354

**Authors:** Gianluca Interlandi

**Affiliations:** Department of Bioengineering, University of Washington, Seattle, WA 98195, USA

**Keywords:** Keywords: Haemostasis and thrombosis, inflammation, methionine oxidation, molecular dynamics simulations, ligand docking, free energy perturbation, anti-thrombotic molecules

## Abstract

The blood protein von Willebrand factor (VWF) is a large multimeric protein that, when activated, binds to blood platelets tethering them to the site of vascular injury initiating blood coagulation. This process is critical for the normal haemostatic response, but especially under inflammatory conditions it is thought to be a major player in pathological thrombus formation. For this reason, VWF has been the target for the development of anti-thrombotic therapeutics. However, it is challenging to prevent pathological thrombus formation while still allowing normal physiological blood coagulation as currently available anti-thrombotic therapeutics are known to cause unwanted bleeding in particular intracranial haemorrhage. This work explores the possibility of inhibiting VWF selectively under the inflammatory conditions present during pathological thrombus formation. In particular, the A2 domain of VWF is known to inhibit the neighboring A1 domain from binding to the platelet surface receptor GpIbα and this auto-inhibitory mechanism has been shown to be removed by oxidizing agents released during inflammation. Hence, finding drug molecules that bind at the interface between A1 and A2 only under oxidizing conditions could restore such auto-inhibitory mechanism. Here, by using a combination of computational docking, molecular dynamics simulations and free energy perturbation calculations, a ligand from the ZINC15 database was identified that binds at the A1A2 interface with the interaction being stronger under oxidizing conditions. The results provide a framework for the discovery of drug molecules that bind to a protein selectively in inflammatory conditions.

## Introduction

The blood protein von Willebrand factor (VWF) has a key function in recruiting blood platelets to the site of vascular injury initiating haemostasis, the physiological process needed to stop blood loss from a wound and promote healing. However, VWF is also thought to have a central role in pathological thrombus formation,^1–3^ and for this reason it has been the target of anti-thrombotic therapies.^4–7^ A challenge in developing anti-thrombotic therapeutics is how to prevent thrombosis while still allowing haemostasis. Currently available anti-thrombotic therapies such as warfarin^8^ or the newly available caplacizumab, which targets VWF directly,^9,10^ increase the risk of haemorrhage, in particular intracranial bleeding. This challenge could be addressed by designing a therapeutic drug that inhibits VWF only when pathological pro-thrombotic conditions are present. It has been shown that an inflammatory state is a major factor contributing to an increased risk of thrombosis.^11^ Inflammation is the natural response in most living organisms to acute events such as traumatic injury or infection. It is accompanied by the release of hydrogen peroxide, which is converted to hypochlorous acid (HOCl) through the action of myeloperoxidase. Such oxidizing agents cause methionine residues in blood proteins to be converted to methionine sulfoxide creating a pro-thrombotic state.^12^ In particular, experimental evidence has shown that the presence of HOCl causes methionine residues in VWF to become oxidized while increasing its platelet-binding function.^13^ Hence, studying the activation mechanism of VWF and how this is altered under oxidizing conditions is essential to find therapeutics that prevent thrombosis while maintaining haemostatis.

The protein VWF has a multimeric structure. Monomers are linked to each other through disulfide bonds at the N- and C-terminii (Figure 1) forming relatively long chains.^14^ Each monomer consists of a number of domains^14^ and of particular importance to the platelet tethering function of VWF are the A1, A2 and A3 domains, which are located adjacent to each other (Figure 1). Under normal conditions, VWF is thought to be coiled up and hence does not significantly bind to blood platelets. However, at the site of vascular injury the A3 domain binds to collagen on the exposed endothelium. Consequently, because of shear in flowing blood the A1 domain becomes exposed and binds to platelet surface receptor glycoprotein Ibα (GpIbα).^15^ Tensile force generated by shear also leads to the unfolding of the A2 domain,^16–19^ which then exposes a scissile bond that is cleaved by the metalloprotease ADAMTS13.^20^ This converts VWF to smaller multimers that are less active in binding platelets. Oxidizing conditions have been shown to have two distinct effects on VWF function. One is that the A2 domain cannot be cleaved by ADMATS13 since a methionine in the cleavage site is converted to methionine sulfoxide in the presence of oxidants.^21^ The other is that VWF is more active in tethering platelets, which is thought to be related to the conversion of methionine residues to methionine sulfoxide in the A domains.^13^ This indicates that under oxidizing conditions VWF undergoes conformational changes that enhance its function. A recent study by the author using a dynamic flow assay indicated that oxidation does not significantly alter the function of the isolated A1 domain but it increases binding between a construct consisting of the A1, A2 and A3 domains to GpIbα.^22^ Also, molecular dynamics (MD) simulations indicated that oxidation of methionines at the interface between A1 and A2 weakens the interaction between the two domains.^22^ This suggests that oxidation removes an auto-inhibitory mechanism in VWF, which is consistent with the observation in previous studies that the A2 domain inhibits A1 from binding to blood platelets.^23,24^ Hence, knowledge of the effects of methionine oxidation may be exploited to develop therapeutics that inactivate VWF selectively under inflammatory conditions. The work presented in this manuscript explores the question whether drug molecules can be designed that bind to oxidized methionine residues at the interface between the A1 and A2 domain significantly stronger than to non-oxidized methionines. Such molecules would then be able to restore the auto-inhibitory mechanism of VWF, which is lost under oxidizing conditions, by stabilizing the A1-A2 complex while having no effect on non-oxidized VWF. For this purpose, the work presented here employs a combination of virtual high-throughput screening using AutoDock Vina, MD simulations and free energy perturbation (FEP) calculations.

**Figure 1:**
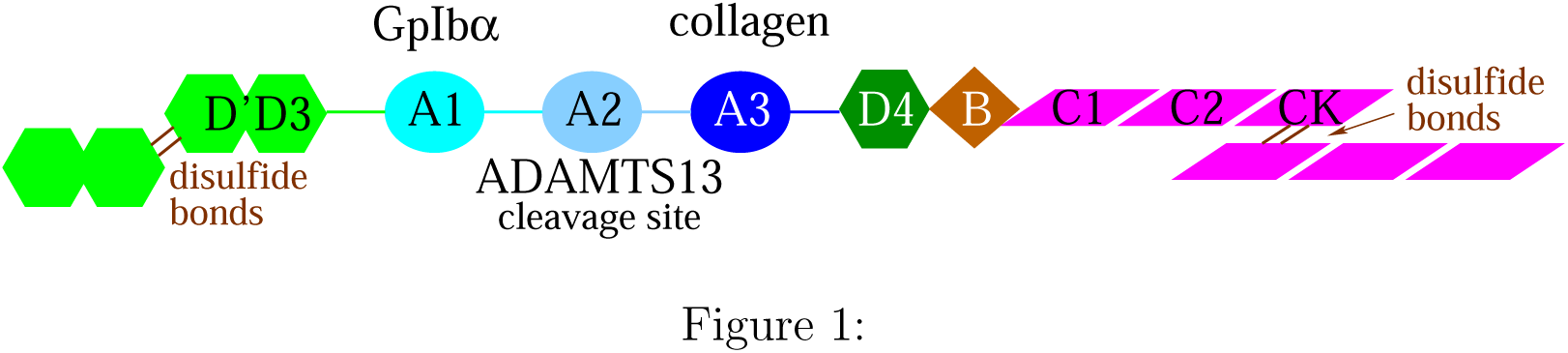
Overview of VWF structure. Schematic representation of a VWF monomer. Indicated are the substrates of the A1 and A3 domain, and the fact that the A2 domain contains the cleavage site for ADAMTS13.

## Materials and Methods

### Initial conformations of the A1A2 complex

Currently, there are no experimentally derived structures of the A1A2 domains complex. Hence, the simulations presented here were performed using models that were obtained in a previous study by others^25^ using a combination of patchdock^26^ and firedock.^27,28^ The coordinates of the models were kindly provided by the authors therein.^25^ In total, there are six different models available, which were used to perform MD simulations in a previous publication by the author.^22^ There, a combination of MD simulations and FEP calculations revealed that oxidation of methionine residues buried at the A1-A2 interface destabilizes the complex between the two domains, which is a possible mechanism for the removal of an auto-inhibitory mechanism and activation of VWF.^22^ Here, four of the six models were selected to use for the screening of possible drug molecules that bind specifically when a methionine residue at the interface is oxidized. Also, for each model a particular methionine site was selected to be targeted by the drug screening. The criteria for selecting the four models and methionine residues were that: 1) the methionine side chain is at the interface but not completely buried, i.e., the solvent accessible surface area (SASA) is larger than 1 Å^2^ as measured along the previously published MD simulations;^22^ and 2) oxidation of the methionine residue has a measurable effect on the free energy of binding between the two domains, i.e., the difference in ΔG between oxidized and non had a p-value < 0.1 (i.e., at least marginally statistically significant) as calculated in a previous publication^22^ (Figures 7 and 6B in reference^22^ present the discussed values for the SASA and ΔΔG, respectively). Using the same labeling as in the previous reference,^22^ these are the combinations of methionine residues and complex models: M1495 in A1A2 1, M1393 in A1A 2, M1495 in A1A2 4, and M1545 in A1A2 6 (Figure 2).

**Figure 2:**
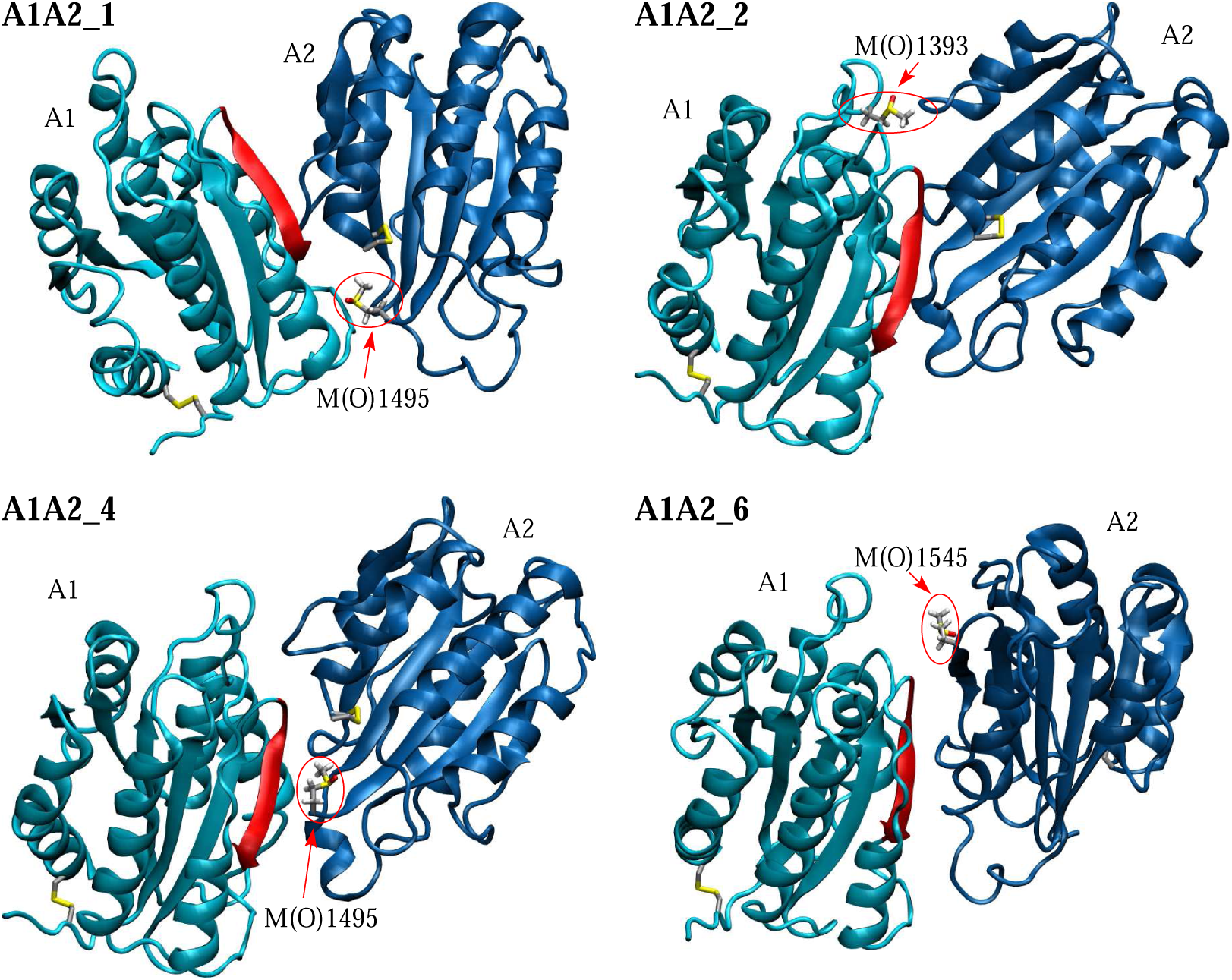
Models of the A1A2 complex and methionine side chains selected to be targeted by drug screening. In the shown structures, the methionine side chains to be targeted have been converted to methionine sulfoxide as described in Section 2 and are labeled and highlighted by red circles. The side chains including cysteines forming disulfide bonds are shown in the stick and ball representation. The backbone of the A1 domain colored in red highlights one of the major contact sites to the platelet surface receptor GpIbα.^57^ The complex models are labeled according to a previous publication.^22^

Prior to performing the virtual screening, the targeted methionine residues were converted to methionine sulfoxide, which consists in the addition of an oxygen atom to the sulphur of the methionine side chain. Then, using the program CHARMM,^29^ 100 steps of steepest descent minimization were performed in vacuo with harmonic constraints applied to all atoms of the complex except the methionine sulfoxide. This was followed by 200 steps of minimization with no constraints applied.

### High-throughput screening with AutoDock Vina

The drug molecules to be tested for binding were downloaded from the ZINC15 database.^30^ The database offers various options to select a subset of drug compounds. For this work, a subset of drug molecules was downloaded using the following two keywords: ”fda”, i.e., approved by the United States Food and Drug Administration (FDA); and ”wait-ok”, i.e., according to the description: ”compounds you can get in 8-10 weeks at modest prices, includes in-stock, agent and on-demand”. The reason behind this selection is to have drug molecules that could be easily obtained and tested later in an *in vitro* assay. Also, by testing drugs approved by the FDA it would be theoretically possible to extend the use of a drug already deemed to be safe to a different purpose such as treating thrombosis. This resulted in a list of in total 1412 unique drug compounds. For several compounds, the list contained multiple possibilities concerning partial charges or the protonation state of non-carbon atoms. Hence, the downloaded list contained a total of 2084 coordinate files. For the purposes of this manuscript, the word ”ligand” refers to a particular coordinate file from the list containing in total 2084 thereof.

The high-throughput virtual screening was performed using the program AutoDock Vina.^31,32^ Prior to the screening, the coordinate files for the ligands (downloaded from the ZINC15 database) and for the receptor (the A1A2 complex model) were converted to the PDBQT format, which specifies which bonds are free to rotate and where only heavy atoms and polar hydrogens are included. The ligands’ coordinate files were converted using the python script prepare ligand4 included in the AutoDock Tools package while the receptor’s coordinate file was converted using the graphical tool ADT. The receptor was generally kept rigid except that bonds in the targeted methionine side chain were free to rotate. A number of bonds in the ligand were also free to rotate as determined by the script prepare ligand4. For the screening, a cubic search box of 15 Å side length was defined centered at the sulphur atom of the targeted methionine side chain. The exhaustiveness parameter was set to 24. In total, four screening runs were performed, one for each of the complex models presented in Figure 2. The computations ran on a six-core Xeon 3.60 GHz processor and lasted each approximately 40 hours.

The docked ligand configurations were then ranked according to the AutoDock Vina score. Only ligands were selected that contained a group able to donate a hydrogen bond so that a stabilizing hydrogen bond between the ligand and the targeted methionine side chain could be formed when the latter is oxidized. For each of the four targeted A1A2 models, at least two ligands were selected corresponding to the highest scores. Only the pose with the highest score was considered for each ligand. Since the scores are rounded to one decimal digit, there can be degeneracy, i.e., multiple ligands can have the exact same score. All ligands corresponding to the highest score were selected. If only one ligand had the highest score, then all ligands corresponding to the second highest score were also selected. The selected model-ligand configurations were then used to start MD simulations (Table 1).

**Table 1:** Simulation Systems. The simulations are grouped by A1A2 model, targeted methionine side chain, and type of simulation. ^a^The name of a simulation is a combination of the A1A2 model, the targeted side chain, the ligand docked, and the type of simulation in the case of FEP. The ligand numbering is according to the list of in total 2084 coordinate files downloaded from the ZINC15 database. The index n is used to denote different replicas (listed in the rightmost column). ^b^Runs to sample the protein-ligand docked state. ^c^Ligands 1340 and 1341 differ in one of the aromatic rings being flipped by 180°. ^d^Ligands 1067 and 1068 differ in the protonation state of one of the nitrogen atoms (doubly protonated in ligand 1067 and single protonated in ligand 1068). ^e^In the FEP simulations, methionine sulfoxide is converted to methionine forward and backward (10 ns each); the runs were started from snapshots sampled in the respective 300-K simulations (40 and 50 ns from replica 1 and 50 ns from replica 2).

| Name<sup>a</sup> | ZINC15 ID | Name | Type | Duration [ns] |
| --- | --- | --- | --- | --- |
| A1A2.1_M(O)1495.1066_n | 3939013 | fosaprepitant | 300 K<sup>b</sup> | 2 x 50 |
| A1A2.1_M(O)1495.1182_n | 64033452 | lumacaftor | 300 K<sup>b</sup> | 2 x 50 |
| A1A2.2_M(O)1393.127_n | 52955754 | ergotamine | 300 K<sup>b</sup> | 15 + 14 |
| A1A2.2_M(O)1393.925_n | 4097286 | budesonide | 300 K<sup>b</sup> | 2 x 50 |
| A1A2.4_M(O)1495.961_n | 1996117 | darifenacin | 300 K<sup>b</sup> | 2 x 50 |
| A1A2.4_M(O)1495.1340_n<sup>c</sup> | 897240<sup>c</sup> | azelastine<sup>c</sup> | 300 K<sup>b</sup> | 2 x 50 |
| A1A2.4_M(O)1495.1341_n<sup>c</sup> | 897240<sup>c</sup> | azelastine<sup>c</sup> | 300 K<sup>b</sup> | 2 x 50 |
| A1A2.6_M(O)1545.360_n | 3831490 | azulfidine | 300 K<sup>b</sup> | 30 + 50 |
| A1A2.6_M(O)1545.1067_n<sup>d</sup> | 1530579<sup>d</sup> | coreg<sup>d</sup> | 300 K<sup>b</sup> | 2 x 50 |
| A1A2.6_M(O)1545.1068_n<sup>d</sup> | 1530579<sup>d</sup> | coreg<sup>d</sup> | 300 K<sup>b</sup> | 2 x 50 |
| A1A2.6_M(O)1545.1180_n | 58581064 | dolutegravir | 300 K<sup>b</sup> | 2 x 50 |
| A1A2.1_M1495.1182_n | 64033452 | lumacaftor | 300 K<sup>b</sup> | 2 x 50 |
| A1A2.2_M1393.925_n | 4097286 | budesonide | 300 K<sup>b</sup> | 20 + 50 |
| A1A2.1_M(O)1495.1182_fep_n | 64033452 | lumacaftor | FEP<sup>e</sup> | 3 x 20 |
| A1A2.2_M(O)1393.925_fep_n | 4097286 | budesonide | FEP<sup>e</sup> | 3 x 20 |

### General setup of the simulations

As stated in reference,^32^ more accurate methods such as MD simulations and free energy perturbation calculations need to be employed in order to study the effects of a mutation on the free energy of biding of a docked ligand. Hence, MD simulations were setup to study the stability of the docked ligands and establish whether a particular ligand binds more strongly to a methionine state in its oxidized than in its unoxidized state.

The MD simulations were performed with the program NAMD^33^ using the CHARMM36m force field^34,35^ in combination with the CHARMM general force field (CGenFF)^36^ and the TIP3P model of water. The force field parameters for methionine sulfoxide were downloaded from the SwissSidechain website^37^ and adapted per analogy to the CHARMM36m force field. The parameters for the ligands were obtained using the CHARMM-GUI online input generator.^38^ Each complex consisting of the A1A2 domains and a docked ligand was inserted into a cubic water box with side length of 100 Å, resulting in a system with in total about 95,000 atoms. The water molecules overlapping with the protein-ligand complex were removed if the distance between any water atom and any atom of the proteins or ligand was smaller than 2.4 Å. Chloride and sodium ions were added to neutralize the system and approximate a salt concentration of 150 mM. To avoid finite size effects, periodic boundary conditions were applied. After solvation, the system underwent 500 steps of minimization while the coordinates of the heavy atoms of the proteins and ligand were held fixed and subsequent 500 steps with no restraints. Each simulation was started with different initial random velocities to ensure that different trajectories were sampled whenever starting with the same initial state. Electrostatic interactions were calculated within a cut-off of 10 Å, while long-range electrostatic effects were taken into account by the Particle Mesh Ewald summation method.^39^ Van der Waals interactions were treated with the use of a switch function starting at 8 Å and turning off at 10 Å. The dynamics were integrated with a time step of 2 fs. The covalent bonds involving hydrogens were rigidly constrained by means of the SHAKE algorithm with a tolerance of 10^−8^. Snapshots were saved every 10 ps for trajectory analysis.

### Equilibration and sampling of the docked state

Before production runs, harmonic constraints were applied to the positions of all heavy atoms of the protein to equilibrate the system at 300 K during a time length of 0.2 ns. After this equilibration phase, the harmonic constraints were released. The systems were simulated for in total a maximum simulation time of 50 ns or the time indicated in Table 1. The first 10 ns of unconstrained simulation time were also considered part of the equilibration and were thus not used for the analysis. During both the equilibration and production phases, the temperature was kept constant at 300 K by using the Langevin thermostat^40^ with a damping coefficient of 1 ps^−1^, while the pressure was held constant at 1 atm by applying a pressure piston.^41^

### Selection of stable protein-ligand complexes

The resulting MD trajectories were analysed to determine stable protein-ligand complexes. A complex between the A1A2 domains and a ligand was considered stable if no detachment was observed in both 50-ns simulations. Detachment was defined if the distance between any atom of the ligand and any atom of the targeted methionine side chain (time-averaged over 4 ns) was larger than 4 Å. Stable protein-ligand complexes where at least one hydrogen bond between the targeted methionine residue and the ligand was observed in at least 10% of the simulation frames of at least one simulation were selected for further analysis (the first 10 of the in total 50-ns long simulations were considered equilibration and excluded from the hydrogen bond analysis).

### Change in free energy of binding upon conversion from methionine sulfoxide to methionine

To determine whether a ligand binds significantly stronger to a methionine when oxidized, the change in the free energy of binding due to the conversion from methionine sulfoxde to methionine was estimated computationally. This was achieved by making use of alchemical transformations^42^ in combination with the thermodynamic cycle^43^ (Figure 3). This is a similar method as in previous publications^22,44^ except that here the methionine side chain in the starting configuration was in its oxidized state as this was used to screen the ligands. The alchemical transformations were performed through free energy perturbation (FEP) calculations.^45^ Each alchemical transformation was performed in the forward and backward direction. In the forward transformation, a methionine sulfoxide side chain is slowly converted to methionine, i.e., the oxygen atom covalently bound to the sulphur atom slowly disappears. The conformation achieved at the end of the forward transformation is then used to start a backward transformation where methionine is converted back to methionine sulfoxide. During the process, the amount of work needed for each transformation is calculated. The forward and backward calculations were then combined and a value for the ΔG of the reduction reaction was obtained using the Bennett’s acceptance ratio method^46^ implemented in the ParseFEP plugin of VMD.^47^ Each forward and backward transformation was performed for 10 ns (20 ns in total) during which a parameter λ was varied from 0 (methionine sulfoxide) to 1 (unoxidized methionine) and from 1 to 0, respectively, in time intervals of the length of 0.1 ns for a total of 100 intermediate states for each direction. The first half of each time window involved equilibration and the second half data collection. A soft core term was introduced to avoid singularities in the van der Waals potential.^48^ To calculate the change in the free energy of binding, ΔΔG, the FEP simulations need to be performed in both, the liganded and the unliganded state. In preliminary simulations with the liganded state, the ligand drifted away from the A1A2 domains as the methionine sulfoxide was transformed to methionine. Hence, harmonic constraints were applied to the carbons of the ligand and the C_α_ atoms of the proteins to prevent the ligand and the A1A2 do-mains from drifting away from each other. The values for the transformation in the unliganded state were derived from previously published simulations performed with just the A1A2 domains.^22^ Each transformation (forward and backward) was performed in triplicates using snapshots sampled along the 50-ns simulations as initial conformations (Table 1), and the results were averaged. By considering the thermodynamic cycle (Figure 3), the difference in the free energy of binding upon the reduction of methionine sulfoxide can be approximated by the difference between the ΔG values calculated from the alchemical transformations in the unliganded and liganded state (see caption of Figure 3).

**Figure 3:**
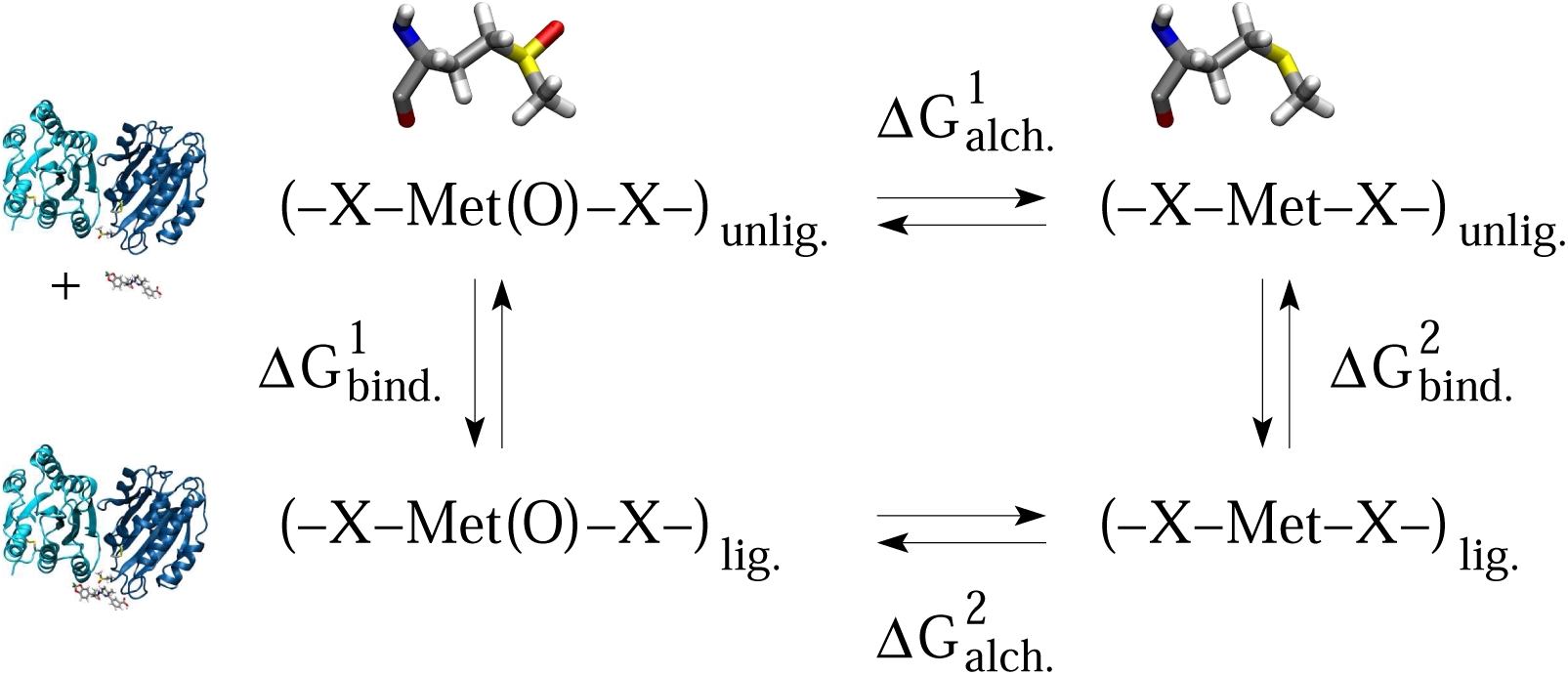
Thermodynamic cycle used to estimate the change in free energy of binding due to the reduction of methionine sulfoxide. The horizontal arrows correspond to the alchemical transformation from methionine sulfoxide to methionine (illustrated at the top) in the unliganded (A1A2 domains) and liganded (A1A2 domains bound to a drug molecule) state (illustrated on the left side). The vertical arrows describe the binding process in the oxidized and unoxidized state. 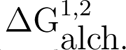 is calculated as described in the text. The change in free energy of binding can then be derived as follows: 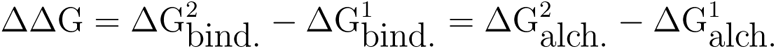

### Determination of persistent contacts and calculation of buried surface area at the protein-ligand interface

The simulation trajectories obtained in the absence of alchemical transformations were screened for the formation of hydrogen bonds between the targeted methionine sulfoxide side chain and the ligand, and also for the formation of side chain contacts between the A1A2 domains and the ligand. To define a hydrogen bond, a H…A distance cutoff of 2.7 Å and a D-H…A angle cutoff of 120° was used, where a donor D could either be an oxygen or a nitrogen, and an acceptor A could be either an oxygen or a nitrogen as long as it is not part of an amino group. A side chain contact between the domains and the ligand was defined to be formed if the distance between any atom of a protein side chain and any atom of the ligand was less than or equal 4 Å. A contact was considered persistent if it was present in at least 66% of the frames of a particular simulation.

The surface area in each individual domain, A1 and A2, buried by the ligand was calculated by subtracting the SASA of each domain determined in the presence of the ligand from the SASA determined in the unliganded state.

## Results

### Screening with AutoDock Vina and stability of the protein-ligand complexes

The program AutoDock Vina^31,32^ offers a simple to use turnkey method to screen drug molecules for binding to a specific site on a protein or protein complex. Here, AutoDock Vina was applied to screen molecules from the ZINC15 database^30^ against the site of methionine residues present at the interface between the VWF A1 and A2 domains and mutated to methionine sulfoxide as expected under inflammatory conditions.^13^ The goal is to find molecules that stabilize the interaction between A1 and A2 selectively under oxidizing conditions. Since currently no experimental structure of the A1A2 domains complex is available, four models of the domains docked to each other were used similarly as in a previous study^22^ (Figure 2, see Section 2 for the criteria how the models and the corresponding methionine residues were selected). In all published models of A1A2, including the four ones used here, the A2 domain obstructs the GpIbα binding site in the A1 domain,^25^ which is consistent with experimental studies showing that the A2 domain has an inhibitory effect on A1.^23,24^ For each of the four complex models, the protein-ligand configurations with the highest AutoDock Vina scores were selected for further analysis (see Section 2 for a description of the criteria how the scores were used to select the liganded configurations). The protein-ligand complexes selected after the AutoDock Vina screening (Table 1) were then used to start MD simulations to study the stability of the obtained liganded configurations and electrostatic contacts between the targeted methionine sulfoxide side chain and the ligand.

For each of the selected protein-ligand complexes, two MD simulations with a total length of 50 ns each were started at 300 K (Table 1). The trajectories were analyzed to determine the stability of the docked ligand conformation and also the presence of hydrogen bonds between the ligand and the targeted methionine sulfoxide side chain. The docked configuration was deemed to be stable if the distance between any atom of the ligand and any atom of the targeted methionine side chain (time-averaged over 4 ns) was less than 4 Å in both simulations (see also Section 2). This was the case in two out of the 11 tested protein-ligand complexes, i.e., model A1A2 1 with lumacaftor (ligand 1182, Figure 5A and Figure 4A) and model A1AA2 2 with budesonide (ligand 925, Figure 5B and Figure 4B, see Supplementary Figures S1-S3 for complexes not deemed to be stable). In both protein-ligand complexes determined to be stable, a hydrogen bond between ligand and methionine sulfoxide side chain was observed to be present in at least one of the two repeat simulations (Figure 4, see Section 2 for details how the presence of hydrogen bonds was determined). The hydrogen bond between the ligand and the sulfinyl oxygen of the methionine sulfoxide side chain is likely to stabilize the ligand in the groove at the interface between the A1 and A2 domain. However, such hydrogen bond would be lost under non-oxidizing conditions thus weakening the protein-ligand interaction.

**Figure 4:**
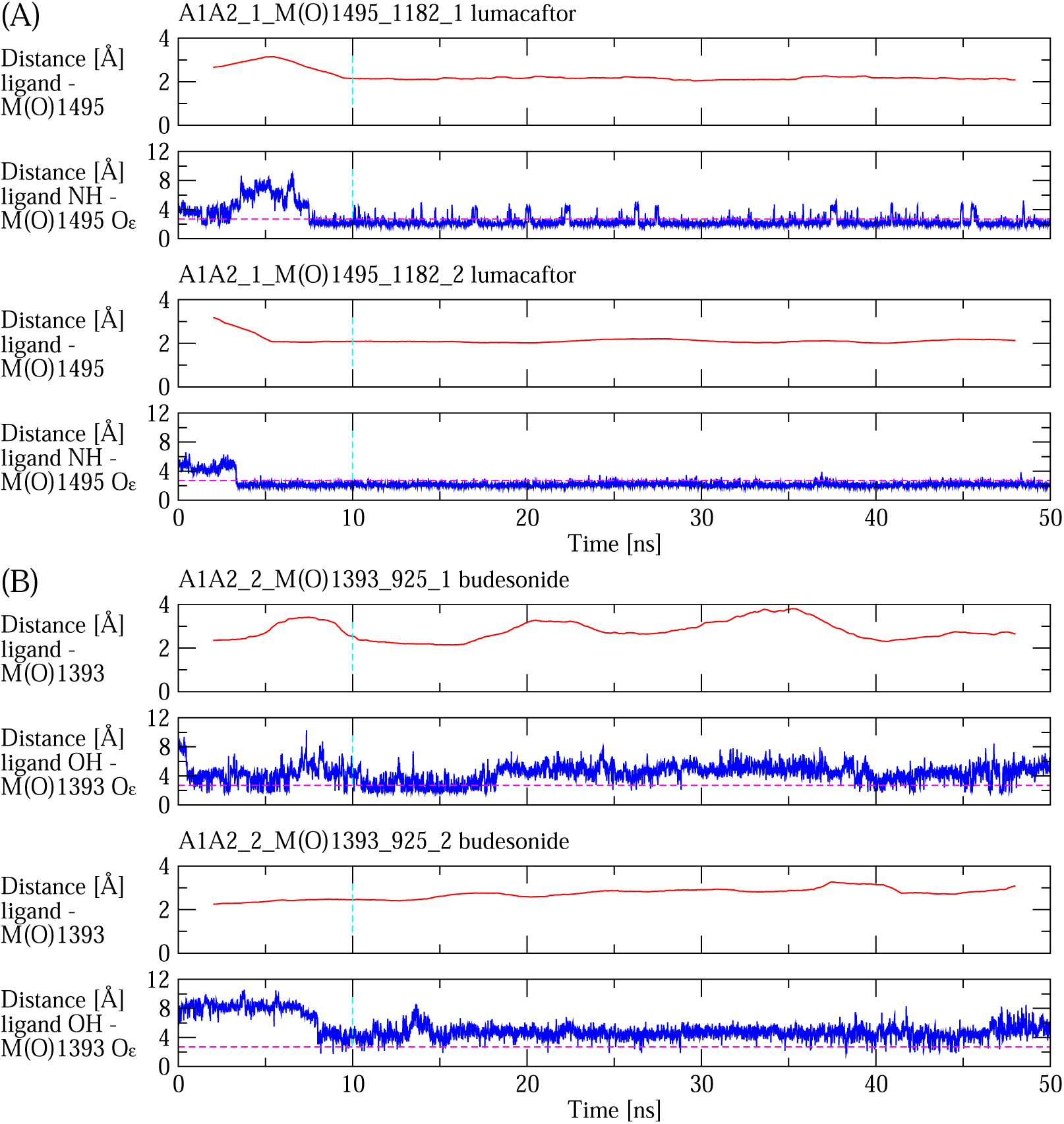
Stability of protein-ligand complexes and hydrogen bond formation. Minimum distance between atoms of the ligand and atoms of the targeted methionine (top) and distance between atoms involved in the hydrogen bond between ligand and methionine sulfoxide side chain (bottom) in simulations with (A) model A1A2 1 and lumacaftor, and (B) model A1A2 2 and budesonide. The minimum distance between ligand and targeted methioinine is averaged over a 4-ns time window. The hydrogen bond, for which the distance is plotted, was found to be formed in at least 10% of the frames in at least one of the two repeat simulations (see also Figure 5). The dashed horizontal magenta line indicates the cutoff of 2.7 Å between the donated hydrogen and the acceptor used to define a hydrogen bond (see Section 2 for a complete description how a hydrogen bond was defined). The dashed vertical cyan line indicates that the first 10 ns of each simulation were considered equilibration and not used to calculate average properties, such as hydrogen bond formation.

**Figure 5:**
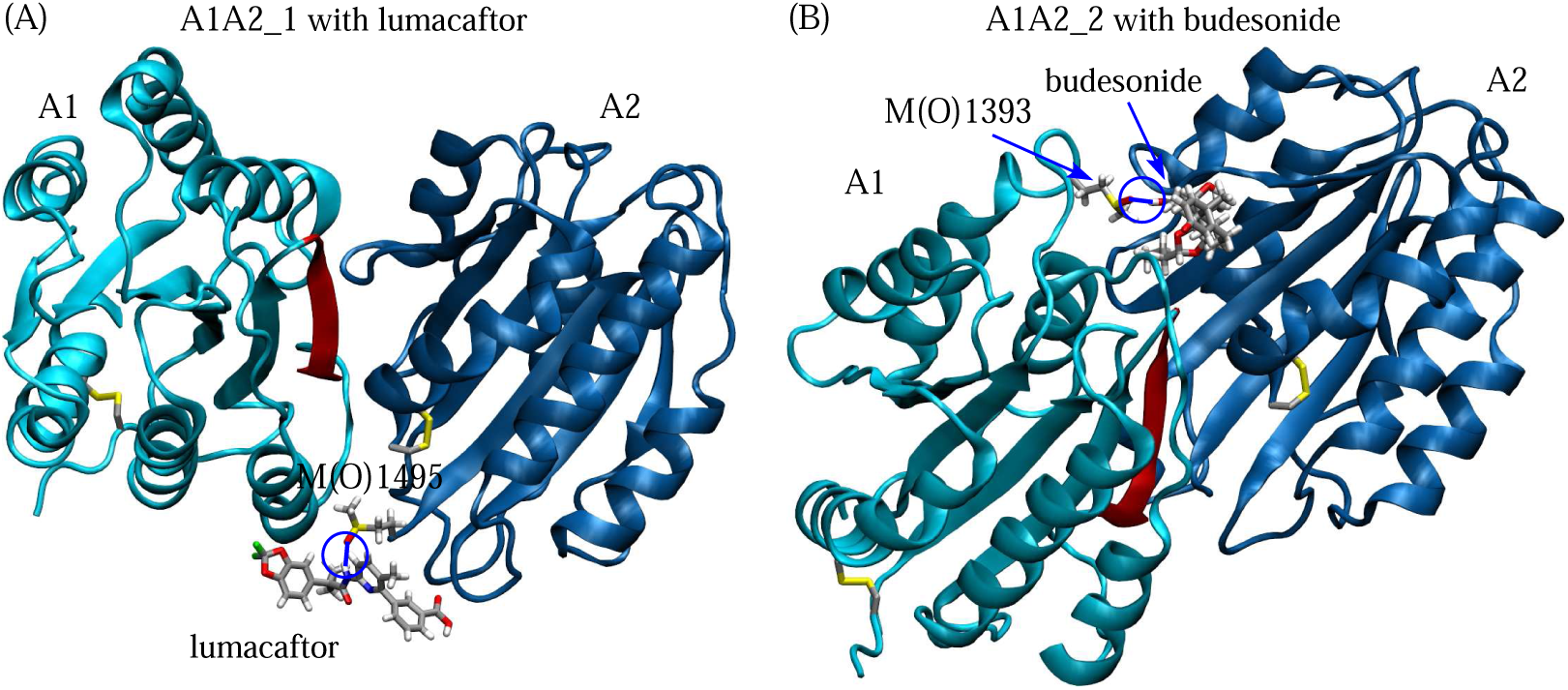
Protein-ligand complexes found to be stable in MD simulations and forming hydrogen bonds between the targeted methionine sulfoxide side chain and the ligand. (A) Lumacaftor docked to model A1A2 1 where M1495 in the A2 domain is oxidized (simulations A1A2 1 M(O)1495 1182 1,2, Table 1). (B) Budesonide docked to model A1A2 2 where M1393 in the A1 domain is oxidized (simulations A1A2 2 M(O)1393 925 1,2, Table 1). The targeted methionine sidechains, disulfide bonds and the ligands are shown in the stick and ball representation. Hydrogen bonds between the respective methionine side chain and the ligand that are present in at least 10% of the frames of at least one simulation are represented by blue lines and highlighted by circles. The backbone of the A1 domain colored in red indicates one of the major contact sites to the platelet surface receptor GpIbα.^57^

### Difference in ligand binding stability between oxidized and non-oxidized methionine through free energy perturbation calculations

After identifying stable protein-ligand complexes (Figure 5), it was necessary to investigate whether a ligand binds to the A1A2 domains specifically when the targeted methionine side chain is in its oxidized state. This means investigating whether binding of the ligand is much weaker when the methionine side chain, which was oxidized when the docking was performed and its stability assessed in MD simulations, loses its sulfinyl oxygen, i.e., it is reduced. Normally, evaluating the change in free energy of binding due to a mutation in the binding pocket requires performing equilibrium binding and unbinding experiments (either *in vitro* or *in silico*) in both, the original state (in this case with methionine sulfoxide) and the mutated state (in this case unoxidized methionine). However, sampling multiple binding and unbinding events in MD simulations is computationally challenging as such events may occur in the microsecond to millisecond time scale.^49^ However, alchemical FEP calculations can significantly reduce the needed computational time. The method used here consists of estimating the work necessary, in this case, to transform methionine sulfoxide to methionine in the liganded and unliganded state and then using the thermodynamic cycle (Figure 3) to estimate the change in the free energy of binding (ΔΔG) between the oxidized and the non-oxidized state of methionine (see Section 2 for details). Similar FEP calculations have been shown to provide values for binding affinities that agree well with estimates from microsecond-long simulations that directly sample multiple binding and unbinding events of small fragments to a protein.^50^

Analysis of the free energy calculations performed here revealed a positive, i.e., unfavorable change in the free energy of binding (ΔΔG) when M(O)1495 was converted to unoxidized methionine in the complex between A1A2 1 and lumacaftor (Figure 6). This indicates that binding of lumacaftor (ligand 1182) is stronger to the oxidized than to the unoxidized methionine at position 1495 of the A2 domain. However, no significant difference was observed between oxidized and unoxidized methionine at position 1393 in the complex between A1A2 2 and budesonide (ligand 925, Figure 6). These results are consistent with the presence of a very stable hydrogen bond between A1A2 1 and lumacaftor (Figure 4A) compared to a weaker hydrogen bond between A1A2 2 and budesonide (Figure 4B). Specifically, the hydrogen bond with lumacaftor (Figure 5A) was present on average in 93% ± 4%, while the hydrogen bond with budesonide (Figure 5B) was observed in 7% ± 5% of the simulation frames averaged each over the last 40 ns of two in total 50-ns long simulations (Figure 4, errors are standard errors of the mean, p-value > 0.05). Loss of such a stable hydrogen bond between A1A2 1 and lumacaftor is likely to be a major contributor to a relatively lower binding affinity when M1495 is unoxidized. Hence, lumacaftor could be a candidate for a molecule that restores binding between A1 and A2, hence inhibiting the platelet-binding function of VWF, specifically under oxidizing conditions.

**Figure 6:**
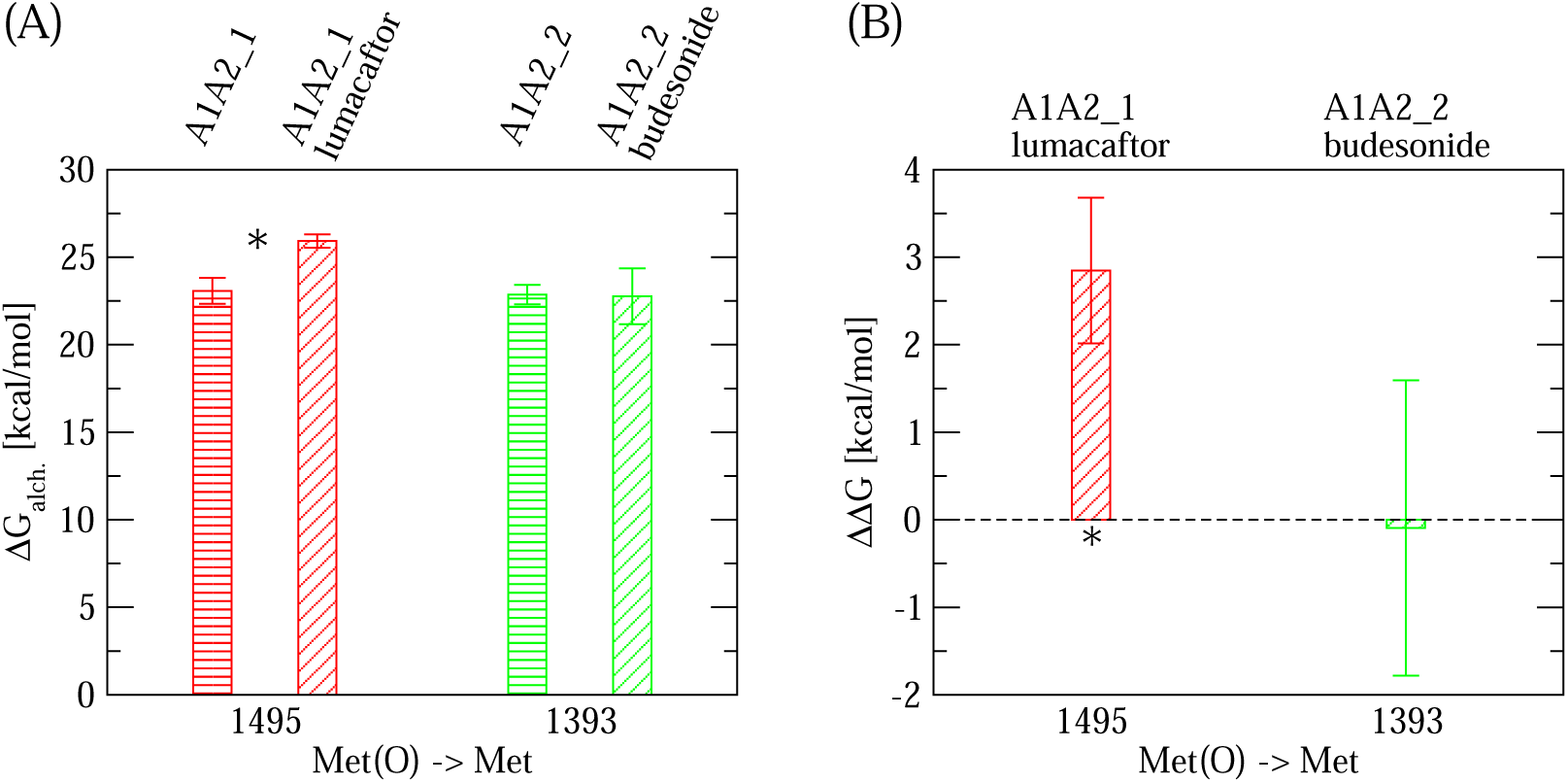
Estimate for the change in free energy of binding upon conversion from methionine sulfoxided to methionine. (A) Calculated ΔG_alch._^1,2^ for the transformation of a given methionine sulfoxide residue to methionine using FEP (see Figure 3 for details). Each transformation was performed in the unliganded A1A2 model (bars with horizontal lines) and in the complex between the respective A1A2 model and the docked ligand (bars with oblique lines). The values for the unliganded simulations were derived from a previous publication.^22^ The structure (isolated A1A2 or A1A2-ligand complex) where the transformation was performed is indicated at the top of the plot. The reported values are averages over three simulations while error bars denote standard errors of the mean. An asterisk (∗) indicates a difference that is statistically significant (p-value calculated from a Student’s t-test smaller than 0.05). (B) The estimated ΔΔG of binding due to the conversion of an individual methionine sulfoxide residue to methionine (see Section 2 and Figure 3). A positive value indicates that reduction of a specific methionine sulfoxide is thermodynamically unfavorable for binding of the respective ligand. Error bars in (B) are derived from the error values displayed in (A) using the error propagation formula: 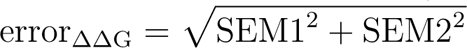 where SEM1 and SEM2 are the standard errors of the mean of ΔG_alch._^1^ and ΔG_alch._^2^, respectively.

Analysis of the SASA of the methionine sulfoxide side chain in the simulations with no FEP (Table 1) revealed that M(O)1495 in the complex with lumacaftor was less exposed than M(O)1393 in the complex with budesonide (Figure 7A) although the difference was not statistically significant (p-value > 0.1). However, the standard deviation of the SASA was much larger for M(O)1393 in the budesonide complex than for M(O)1495 in the lumacaftor complex (Figure 7A, p-value < 0.05) indicating that the latter was fluctuating less. Furthermore, the donor group in the ligand that was involved in the hydrogen bond with the A1A2 domains was much more protected from the solvent in the complex with lumacaftor than in the one with budesonide (Figure 7A, p-value < 0.05). Since the methionine sulfoxide side chain is hydrophilic (because of the presence of a sulfinyl group) its sequestration from solvent would be energetically unfavorable. However, such an energetic penalty is probably compensated by the formation of a stable hydrogen bond between M(O)1495 and lumacaftor. Buried hydrogen bonds are thought to contribute to the thermodynamic stability of a folded protein, the complex between two proteins, and in particular the complex between a protein and a ligand.^51^ Thus, the hydrogen bond between M(O)1495 and lumacaftor is likely to significantly stabilize the protein-ligand complex and its loss due to the reduction of the methionine results in the lower binding affinity between A1A2 and the ligand (Figure 6B). Hence, the change in the ΔG of binding can be said to be correlated to how well the protein-ligand hydrogen bond is protected from the solvent and how stable it is, as evidenced by the comparison between the lumacaftor and budesonide complex (Figure 6B).

**Figure 7:**
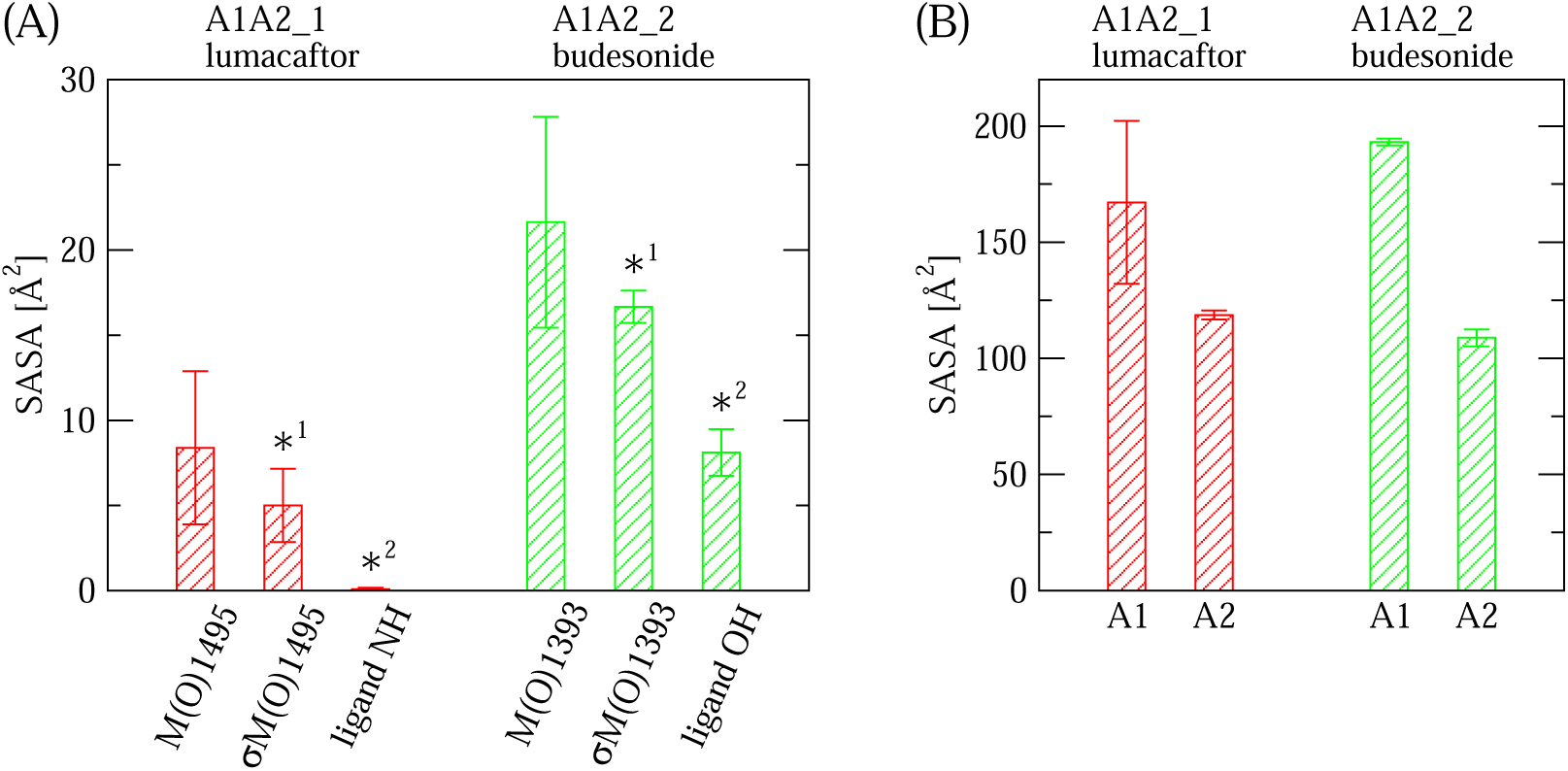
Surface area analysis of the protein-ligand binding interface. (A) Solvent accessible surface area (SASA) of the targeted methionine side chain (M(O)1495 in A1A2 1 with lumacaftor and M(O)1393 in A1A2 2 with budesonide) and the donor group in the ligand (NH in lumacaftor and OH in budesonide), which are involved in hydrogen bond formation (Figures 4-5). σM(O)1495 and σM(O)1393 refer to the standard deviations of the SASA along trajectories. The reported values are averages over the last 40 ns of two in total 50-ns long simulations with each respective complex (A1A2 1 M(O)1495 1182 1,2 and A1A2 2 M(O)1393 925 1,2, Table 1). Error bars are standard errors of the mean. ∗^1^ and ∗^2^ denote that the differences between the corresponding values for the two complexes are statistically significant (p-value < 0.05 from a Student’s t-test). (B) Surface area of each domain buried by the ligand. Reported are averages over two simulations and standard errors of the mean.

Analysis of the buried SASA between A1A2 and the ligand revealed that in both, the lumacaftor and the budesonide complex, the ligand shields part of the surface area of both, the A1 and A2 domain from the solvent (Figure 7B, see Section 2 for details about the calculation). Burial of surface area normally accessible to solvent in the unliganded state indicates that the ligand contacts both domains. This is necessary for a drug molecule to restore the inhibitory function that the A2 domain normally has on the A1 domain, i.e., the ligand acts as a glue to extend the contact surface between the two domains. However, in the case of lumacaftor oxidation of M1495 acts as a switch whereby the ligand binds significantly stronger under oxidizing conditions, which is a desired property for a drug molecule that inhibits the pathological thrombotic action of VWF while allowing its physiological haemostatic function.

### Analysis of contacts between A1A2 and lumacaftor

The complex between A1A2 1 and lumacaftor provides a framework for the design of drug molecules that restore the inhibitory function of A2 onto the A1 domain selectively under oxidizing conditions. Such a molecule needs to contact both the A1 and A2 domain in order to provide an extension of the inter-domain binding interface and enhance the binding affinity. Hence, the trajectories A1A2 1 M(O)1495 1182 1,2 (Table 1) were screened for persistent contacts between lumacaftor and side chains of the A1A2 domains.

A contact between a side chain and an atom of the ligand was defined to be formed when at least one atom of the side chain was within 4 Å of the particular atom of the ligand. A contact was determined to be persistent if it was present in at least 66% of the simulation frames in at least one of two simulations with the complex. Side chains of A1A2 and atoms of lumacaftor that were involved in at least one persistent contact are illustrated in Figure 8. In both simulations, R1336, S1338, E1339, R1341, R1342 and M(O)1495 were involved in persistent contacts highlighting that lumacaftor contacts both the A1 and A2 domain (Figure 8). The molecule lumacaftor consists of three major aromatic rings. All atoms of the ring containing two fluoride atoms and most atoms of the other two rings are involved in persistent contacts (Figure 8). These observations suggest that aromatic rings provide stabilizing hydrophobic contacts between the ligand and the A1A2 domains and may be an important consideration in the design of a molecule meant to bind at the inter-domain interface. At the same time, the amide group situated between two of the rings (Figure 8) provides the ability to form a stable hydrogen bond between the methionine sulfoxide side chain and lumacaftor, which significantly strengthens the protein-ligand binding selectively under oxidizing conditions.

**Figure 8:**
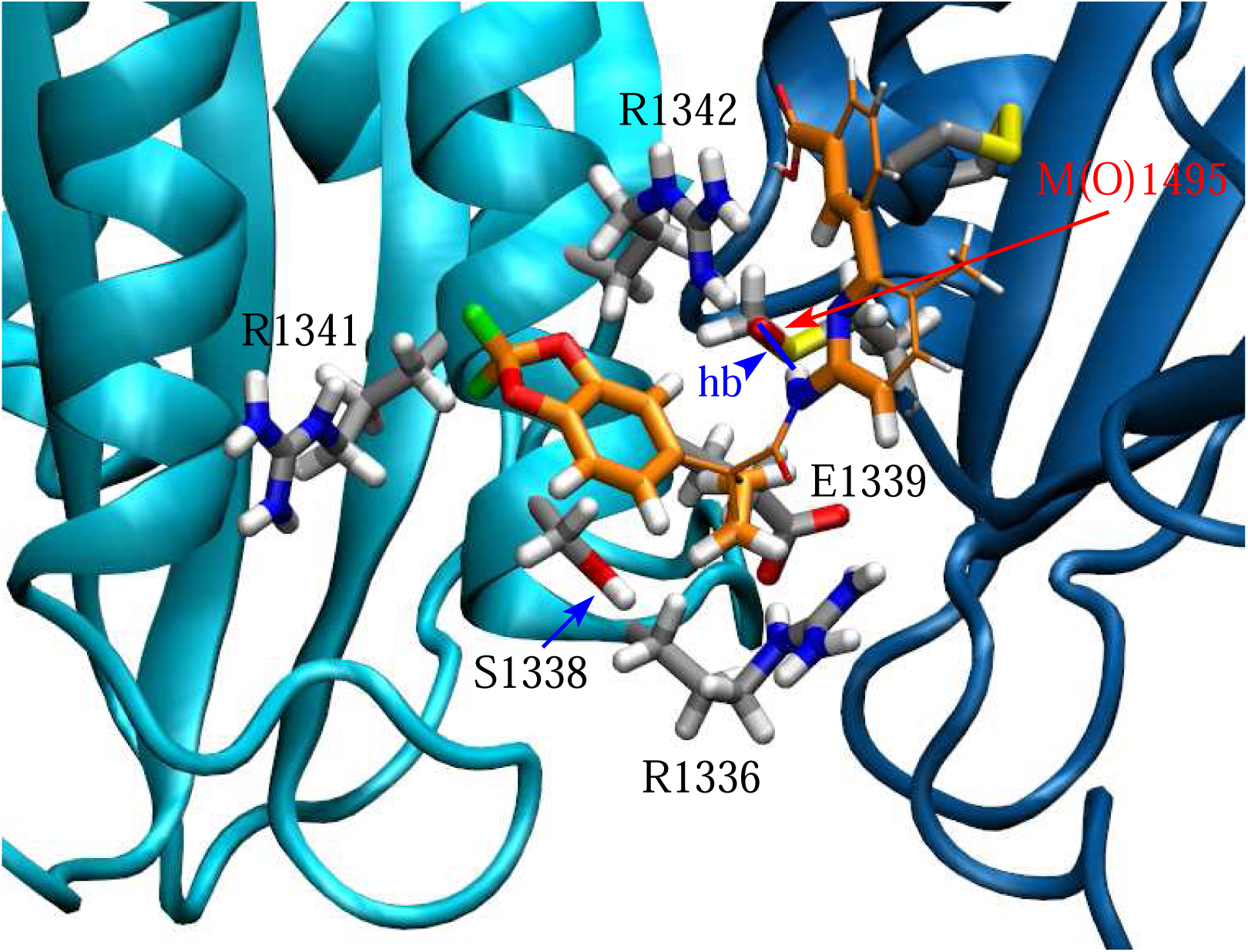
Side chains of A1A2 and atoms of lumacaftor involved in persistent contacts. Side chains that were involved in persistent contacts with the ligand in the simulations A1A2 1 M(O)1495 1182 1,2 (Table 1, see Section 2 for definition of persistent contacts) are displayed in the stick and ball representation and labeled. The ligand is also displayed in the stick and ball representation, where atoms that were not involved in persistent contacts with protein side chains have bonds represented with a smaller radius. To distinguish side chains from the ligand the carbons of the latter are colored in orange. The hydrogen bond (hb) observed in the simulations between the side chain of M(O)1495 and lumacaftor is highlighted by a blue line. The cysteine bond in the A2 domain is also shown for the sake of completeness although it was not involved in contacts with the ligand.

## Discussion

A major challenge in the design of anti-thrombotic therapeutics is how to prevent the pathological formation of a thrombus while at the same time maintaining the physiological ability of blood to coagulate at the site of injury. Inflammation is a major driver of thrombotic events^11^ and oxidizing agents released during the inflammatory response alter the function of blood proteins by converting methionine residues to methionine sulfoxide.^12^ In VWF, oxidation of methionine residues in the A1, A2 and A3 domains has been linked to an increased binding activity to platelets, and VWF is known to play a central role in pathological thrombus formation.^13^ However, inhibiting VWF may disrupt its normal haemostatic function leading to unwanted bleeding. The importance of VWF in the physiological process of haemostasis is highlighted by the fact that mutations in VWF lead to von Willebrand disease, a minor bleeding disorder.^14,44,52^

This manuscript proposes developing a therapeutic that inhibits VWF selectively when oxidation-induced thrombotic conditions are present. The protein VWF is known to have an auto-inhibitory mechanism whereby the function of the A1 domain, which binds to the platelet surface receptor GpIbα, is inhibited by neighboring regions.^23,24,53–55^ This inhibitory mechanism is removed either under tensile force, such as generated in rapidly flowing blood,^15,56,57^ or in inflammatory conditions through oxidation of methionine residues.^13^ Finding a drug molecule that restores such auto-inhibitory mechanism selectively in the presence of oxidized methionine residues would provide a way to block VWF function where it could lead to pathological thrombus formation but not where it is needed for its normal physiological activity. In particular, the A2 domain has been shown to reduce binding of VWF to blood platelets,^23,24^ which could be due to direct obstruction of the GpIbα-binding site in A1^25^ (Figure 2). A previous analysis using models of the A1A2 domains^25^ indicated that oxidation of methionine residues located at the inter-domain interface destabilize the complex between the two proteins.^22^ Hence, a virtual high-throughput screening method was used here to search for molecules that bind at the site of methionine residues located at the inter-domain interface when such methionines are in their oxidized state. The protein-ligand complexes with the highest energy scores were then tested in MD simulations and those that were found to be stable and presented a hydrogen bond between the methionine sulfoxide side chain and the ligand were selected for further analysis. This yielded two complexes, one between the model A1A2 1 (containing M(O)1495) and lumacaftor and one between A1A2 2 (containing M(O)1393) and budesonide (Figure 5). The desired property of such a molecule is that it binds under oxidizing conditions much more strongly than under normal non-oxidizing conditions. Hence, FEP calculations coupled with the thermodynamic cycle were performed to evaluate the difference in free energy of binding between the targeted methionine residue in its oxidized and in its non-oxidized state. The analysis revealed that this was the case for the complex between A1A2 1 (containing M(O)1495) and lumacaftor but no statistically significant difference was found for the complex between A1A2 2 (containing M(O)1393) and budesonide (Figure 6). A major reason for this difference is probably the fact that the hydrogen bond involving methionine sulfoxide was much more stable in A1A2 1-lumacaftor than in A1A2 2-budesonide (Figure 4). Furthermore, the SASA of M(O)1495 in A1A2 1 was found to fluctuate much less than M(O)1393 in A1A2 2 and the donor group for the hydrogen bond was much more buried in lumacaftor than in budesonide (Figure 7A). Buried hydrogen bonds have been found to significantly contribute to the thermodynamic stability of protein-ligand complexes and to be important for molecular recognition.^51^ Taken together, the change in free energy of binding when comparing oxidized to non-oxidized methionine in the complex between A1A2 1 and lumacaftor is probably accounted for by the loss of the protein-ligand hydrogen bond. Hence, the hydrogen bond between A1A2 1 M(O)1495 and lumacaftor provides specificity whereby binding is weakened upon loss of methionine oxidation.

In both complexes, A1A2 1-lumacaftor and A1A2 2-budesonide the ligand contacts both domains (Figure 7B) indicating that the contact area between the two domains is expanded by the ligand docking. Hence, the drug molecule acts as a ”glue” further enhancing the inter-domain interaction. However, in A1A2 1-lumacaftor the interaction is specific to M(O)1495 being in its oxidized state. This makes lumacaftor a candidate for a drug that restores the inhibitory function of the A2 domain specifically under oxidizing conditions. Notably, the A1A2 1-lumacaftor can serve as a framework for the design of further drug molecules with similar specific binding properties that can be tested in a laboratory setting. Analysis of the interface between the domains and the ligand revealed that aromatic rings provide stabilizing hydrophobic contacts (Figure 8) and hence this is an important design consideration.

Four conclusions emerge from this work. The first is that a combination of virtual high-throughput docking, MD simulations and alchemical transformations (here performed through FEP) conbined with the thermodynamic cycle can be used to find drug molecules that are specific to a post-translational modification in proteins such as methionine oxidation. The program AutoDock Vina^31,32^ is a powerful tool for the rapid search of drug molecules that bind to a specific target site of a protein. However, subtle changes in the protein primary sequence due to a mutation or a post-translational modification require combining the results of the *in silico* docking with MD simulations and alchemical free energy calculations.

The second conclusion is that a buried hydrogen bond between the sulfinyl group of the methionine sulfoxide side chain and the ligand provides specificity for the molecular interaction. This is consistent with a study analysing protein-ligand interactions in the protein data base (PDB), which found that hydrogen bonds provide specificity although it did not include oxidized methionine side chains.^51^ The study presented here provides evidence that hydrogen bonds involving a methionine sulfoxide side chain also provide specificity in protein-ligand recognition.

The third conclusion is that methionine oxidation, which turns the side chain from hydrophobic to hydrophilic, acts as a switch for the binding of lumacaftor to the A1A2 domains. Hence, lumacaftor, or any drug with similar properties, would bind specifically under oxidizing conditions generated by a pro-thrombotic inflammatory state. Since the ligand binds to both, the A1 and A2 domain, the inter-domain interaction is strengthened out-weighing the disruptive effect of methionine oxidation and hence restoring the inhibitory function of the A2 domain onto A1. This would then inhibit VWF from binding platelets in an inflammatory pro-thrombotic environment. However, in the absence of oxidation lumacaftor, or a molecule with similar properties, would bind weakly so that VWF can exert its physiologic function in preventing blood loss at the site of injury.

The fourth conclusion is that thanks to computational atomistic modeling it is possible to analyse what interactions are important between a targeted protein or protein complex and a ligand especially when such ligand needs to bind according to a specific post-translational modification. Here, by analysing the A1A2 1-lumacaftor interface it was possible to recognize that, besides the hydrogen bond between M(O)1495 and the ligand, hydrophobic interactions between aromatic rings of the ligand and the hydrophobic parts of side chains are key to stabilize the interaction. Hence, the complex between A1A2 and lumacaftor studied here can be used as a model framework to design further examples of molecules with similar properties such that they bind specifically in the presence of methionine sulfoxide. Molecules with such properties can then be tested *in vitro* in functional assays.

In conclusion, this study provides a method to identify *in silico* a molecule that binds at the interface between the A1 and A2 domain of VWF selectively when a methionine residue is oxidized. The modeled protein-ligand complex provides insights into key contacts between the drug molecule and the A1A2 domains such as a buried hydrogen bond involving the targeted methionine sulfoxide side chain and hydrophobic contacts with aromatic rings of the ligand. Such a therapeutic molecule can then restore the auto-inhibitory mechanism of VWF selectively under oxidizing conditions that are present during an inflammatory state. The method used here may also be applied to other blood proteins whose function is also regulated by methionine oxidation.^12^ The overall goal is the discovery of a therapeutic that blocks the pathological thrombus formation that leads to thrombotic events while maintaining the physiological haemostatic response that prevents blood loss at the site of vascular injury.

## Acknowledgments

I would like to thank Marino Convertino for helpful discussions about molecular docking. The simulations were performed on the Expanse supercomputer at the San Diego Supercomputing Center thanks to a XSEDE allocation^58^ with grant number TG-MCB140143, which is made available through support from the National Science Foundation. This research was financially supported by NIH grant R01HL153253 and a Phase 1 Technology Commercialization Award from the Washington Research Foundation.

