## Supplemental data for "Exploring ligands that target von Willebrand factor selectively under oxidizing conditions through docking and molecular dynamics simulations"

**Gianluca Interlandi**

Supplemental data

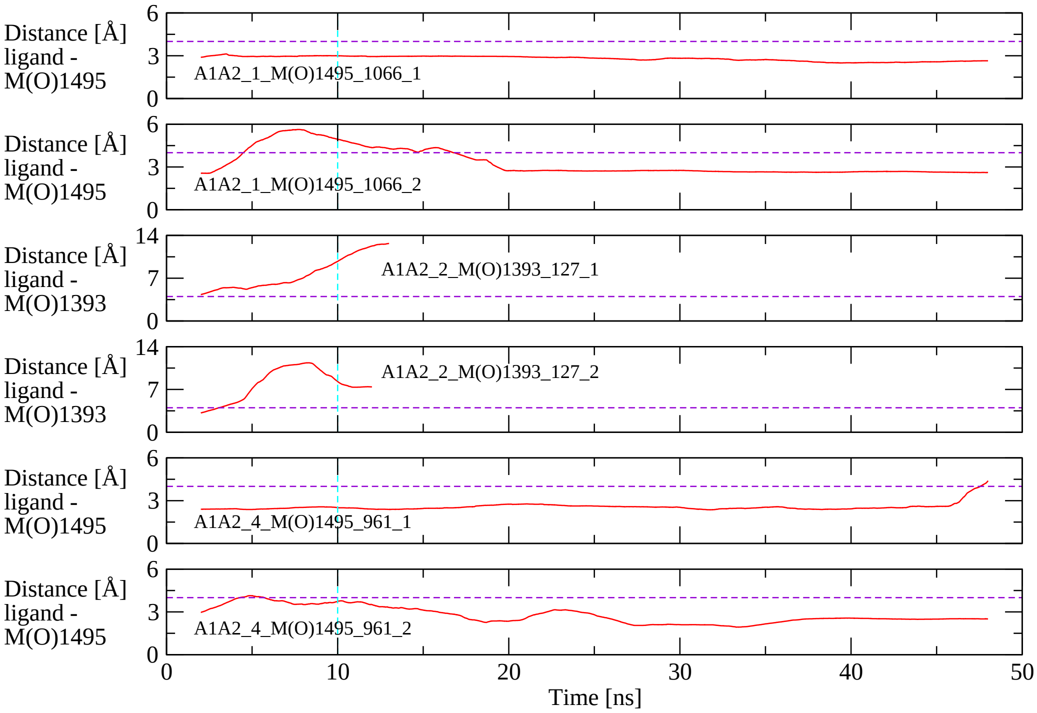


Figure S1: **Stability of complexes between the A1A2 domains and ligands 1066, 127 and 961.** Shown is the 4-ns time averaged minimum distance between atoms of the ligand and atoms of the targeted methionine side chain in simulations with the complex between A1A2 and ligands. Two simulations were performed with each complex. The dashed horizontal violet line indicates the cutoff of 4 Å above which the domains-ligand complex is not considered stable. The dashed vertical cyan line indicates that the first 10 ns of each simulation were considered equilibration and not used to calculate average properties, such as hydrogen bond formation.

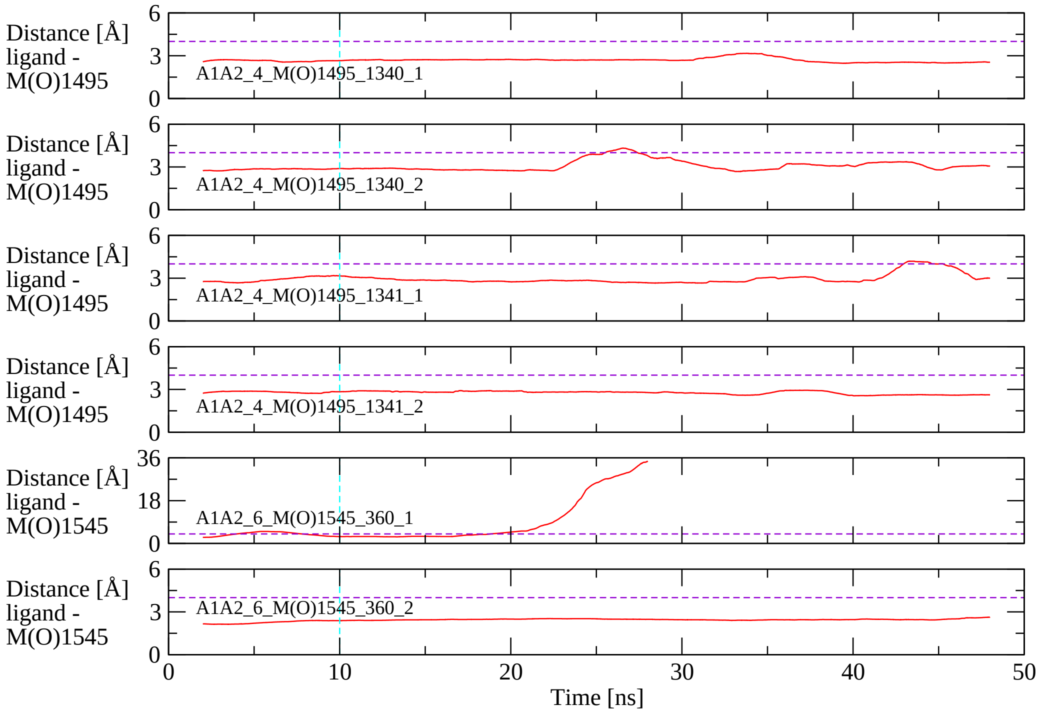


Figure S2: **Stability of complexes between the A1A2 domains and ligands 1340, 1341 and 360.** Shown is the 4-ns time averaged minimum distance between atoms of the ligand and atoms of the targeted methionine side chain in simulations with the complex between A1A2 and ligands. Two simulations were performed with each complex. The dashed horizontal violet line indicates the cutoff of 4 Å above which the domains-ligand complex is not considered stable. The dashed vertical cyan line indicates that the first 10 ns of each simulation were considered equilibration and not used to calculate average properties, such as hydrogen bond formation.

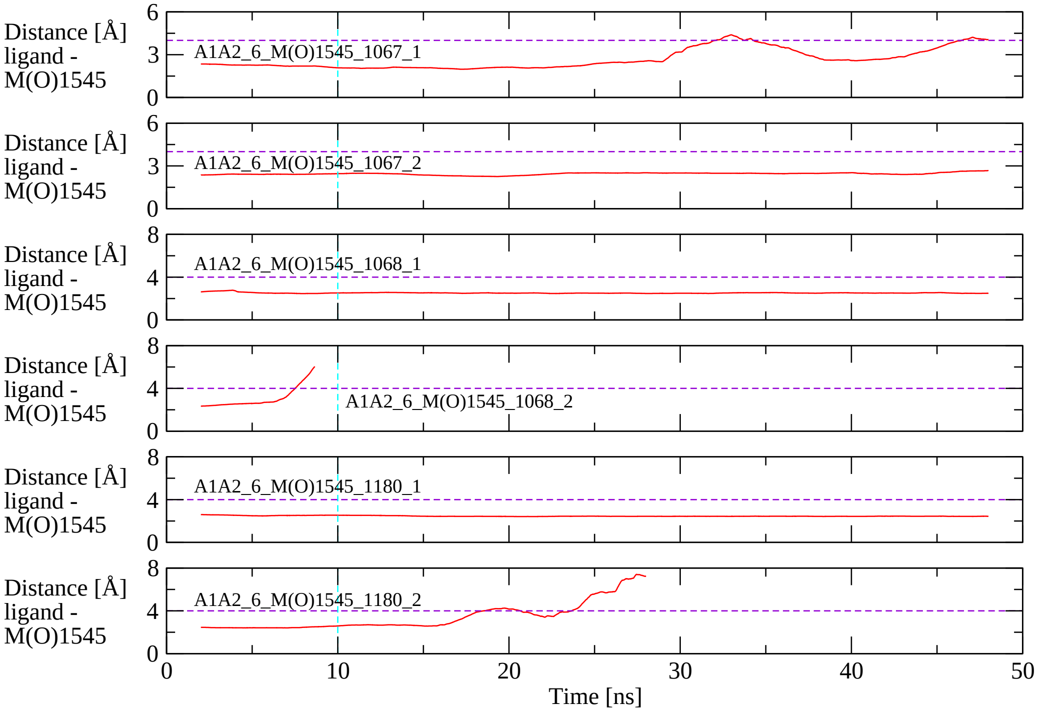


Figure S3: **Stability of complexes between the A1A2 domains and ligands 1067, 1068 and 1180.** Shown is the 4-ns time averaged minimum distance between atoms of the ligand and atoms of the targeted methionine side chain in simulations with the complex between A1A2 and ligands. Two simulations were performed with each complex. The dashed horizontal violet line indicates the cutoff of 4 Å above which the domains-ligand complex is not considered stable. The dashed vertical cyan line indicates that the first 10 ns of each simulation were considered equilibration and not used to calculate average properties, such as hydrogen bond formation.
